# Impact of mothers’ past experience and early-life stress on aggression and cognition in adult male mice

**DOI:** 10.1101/453084

**Authors:** V. Reshetnikov, Yu. Ryabushkina, N. Bondar

## Abstract

Early life is an important period for brain development and behavioral programming. Both reduced maternal care and stress in early life are risk factors for various psychiatric disorders. Here, we hypothesized that females’ stressful experience in their early life can lead to a disruption of mother-offspring interactions toward their own progeny. The objective of this study is to assess the effects of mothers’ past stressful experience, early-life stress alone or both on behavior in adult male mice. In this study, female mice were allowed to raise their pups either without exposure to stress (normal rearing condition, NC) or with exposure to maternal separation (3h/day, maternal separation, MS) on postnatal days 2–14. Adult F1 female mice who had experienced MS (stressed mothers, SM) or had been reared normally (undisturbed mothers, UM) were used for generating F2 offspring to be or not to be further exposed to early-life stress. We assessed anxiety-like behavior, exploratory activity, locomotor activity, aggression and cognition in four groups of adult F2 males (UM+NC, UM+MS, SM+NC, SM+MS). We found that SM+MS males become more aggressive if agonistic contact is long enough, suggesting a change in their social coping strategy. Moreover, these aggressive males tended to improve longterm spatial memory. Aggressive SM+NC males, in contrast, showed learning impairments. We did not find any significant differences in anxiety-like behavior or exploratory and locomotor activity. Overall, our findings suggest that mothers’ early-life experience may have important implications for the adult behavior of their offspring.

## 1. Introduction

Early postnatal periods are utterly important for individual development and shaping the behavioral phenotype of adults (Teicher et al., 2016, Krugers et al., 2017, Maccari et al., 2017). Early-life stress as well as low levels of maternal care may lead to behavioral impairments and cognitive deficits in adulthood (Champagne et al., 2003, Pryce and Feldon, 2003, Kaffman and Meaney, 2007, Kosten et al., 2012). Rodent studies showed that untoward events in early life lead to enhanced anxiety and impair learning and memory (Kosten et al., 2012, Loi et al., 2017). Human studies also demonstrate that stress and maltreatment in childhood may increase the risk of psychic and behavioral impairments and lead to enhanced anxiety, aggression and impulsivity in adulthood (Johnson et al., 2002, Chen et al., 2012, Vaiserman, 2015, Chistiakov and Chekhonin, 2017).

Awareness is growing that, apart from delayed adverse effects on adult behavior, early postnatal stress lead to consequences that have effects on the behavior of the progeny in at least two or more consecutive generations (Franklin et al., 2011, Babb et al., 2014, Schmauss et al., 2014, van Steenwyk et al., 2018). Effects of early postnatal stress may be passed down through generations via both germ line-dependent (Crews et al., 2012) and non-genomic mechanisms, with epigenetic factors of shaping the behavioral phenotypes of the progeny (Francis et al., 1999, Fleming et al., 2002, Champagne et al., 2003, Weaver, 2007, van Steenwyk et al., 2018). Different levels of mother-offspring interaction during early postnatal periods are associated with changes in DNA methylation of promoter elements and offspring behavior (Kaffman and Meaney, 2007). Disruptions in these interactions observed, for example, in females exposed to various types of stress in their early life may have consistent consequences passed down through generations (Babb et al., 2014, Schmauss et al., 2014, Nephew et al., 2017). Nevertheless, although it is deemed to be obviously important to study the effects of females’ past experience on their future interactions with their pups, with implications for the pups in adulthood, the number of such studies is limited. Some studies in rats proved that prolonged maternal separation reduces maternal licking and crouching over pups in adult F1 females (Lovic et al., 2001), while early-life handling significantly increases the frequency of maternal licking/grooming and arched-back nursing in the female offspring of low-licking mothers, but has no effect on the offspring of high-licking mothers (Francis et al., 1999). Our results also demonstrated that adult females, who had been exposed to prolonged maternal separation in their early life showed a reduced licking level to their own progeny (in press). In addition to the maternal separation model, the main aspect whereof is tactile isolation (Kaffman and Meaney, 2007), B. Nephew and C. Murgatroyd with the co-workers scrutinized the effects of social stress (having to be next to an intruder male for an hour once a day) in early life on maternal care in adult females (Murgatroyd and Nephew, 2013). Their results demonstrate that the effects of chronic social stress during lactation adversely affect maternal care in and the hormonal state of lactating F0 mothers, as well as maternal care given by adult F1 females to their progeny (Carini and Nephew, 2013, Murgatroyd and Nephew, 2013). Moreover, it was found that the consequences of reduced maternal care (F0 and F1) have implications for the next generation (F2) and lead to reduced social behavior in F2 juveniles and adults (Babb et al., 2014). At the same time, the only relevant study in mice known to us demonstrated that prolonged maternal separation in combination with additional stress (20-min restraint in a Plexiglas tube or 5-min forced swimming in cold water) applied to F0 mothers did not lead to significant changes in the adult F1 females’ maternal care; however, the behavior of their adult F2 progeny was changed (Weiss et al., 2011). Thus, this list of evidence lends further support to the hypothesis that the effects of stress in early life may have implication for several generations.

However, although rodent studies provide evidence that mothers’ past experience as well as early-life stress in pups may be reasons behind behavioral and hormonal impairments in adult progeny, their contributions to shaping behavioral phenotypes are not yet compared. In this work, we proposed that mothers’ past experience in their early life may affect their interactions with their own pups, which may be an additional and probably more important risk factor for the development of behavioral deficiencies in the adult offspring than these own stress in their early life. The aim of our study was to assess the effects of mothers’ past experience in their early life, the effects of stress in their pups due to prolonged maternal separation in their early life, and the effects of these stresses in combination on the behavior of the adult male mouse offspring. To assess various behavioral patterns, we made use of a test battery, which evaluates the parameters of individual and social behavior, aggression, learning, and behavioral effects of prolonged social isolation.

## 2. Methods

### 2.1. Animals

C57BL/6 mice were maintained at the Animal Facility of the Institute of Cytology and Genetics of the Siberian Branch of the Russian Academy of Sciences (RFMEFI62117X0015) (Novosibirsk, Russia). All procedures were approved by the Ethical Committee of the Institute of Cytology and Genetics, SB RAS, (Protocol #25, December 2014). The animals were housed under standard conditions (a 12 × 12 h light/dark cycle, lights on at 8:00 a.m.; food (pellets) and water available *ad libitum*).

### 2.2. Maternal separation

As a model of early-life stress, we used the repeated dam–offspring separation described previously (Bondar et al., 2018b, Reshetnikov et al., 2018b). Briefly, pups were separated from their mothers for 3 h once a day during the first two weeks of life. The pups were removed from their home cage and placed individually into small boxes filled with bedding. During separation, the temperature was kept at 30 ± 2°C using a heat mat to prevent thermoregulatory distress. The control pups were not separated from their dams. On postnatal day 30, all litters were weaned and the young animals remained together with their same-sex littermates under standard housing conditions until adulthood.

### 2.3. Experimental design

Twenty-four adult female (F0) and 9 males were used for generating F1 offspring (Figure 1). Each pregnant female was placed alone in a new cage at least two days prior to parturition. F1 pups were exposed to prolonged maternal separation (see section 2.2.) or reared under normal conditions. Then, two groups of adult F1 females (stressed mothers, SM, and unstressed mothers, UM) 2.5-3 months of age were mated with naive males for generating F2 pups. F2 pups from each group of females were exposed to prolonged maternal separation or reared under normal conditions. Thus, the following four groups of F2 offspring were used in the experiment:

1. Unstressed mother + pups, normal rearing condition, UM+NC
2. Stressed mother + pups, normal rearing condition, SM+NC
3. Unstressed mother + maternal separation of pups, UM+MS
4. Stressed mother + maternal separation of pups, SM+MS

**Figure 1.**
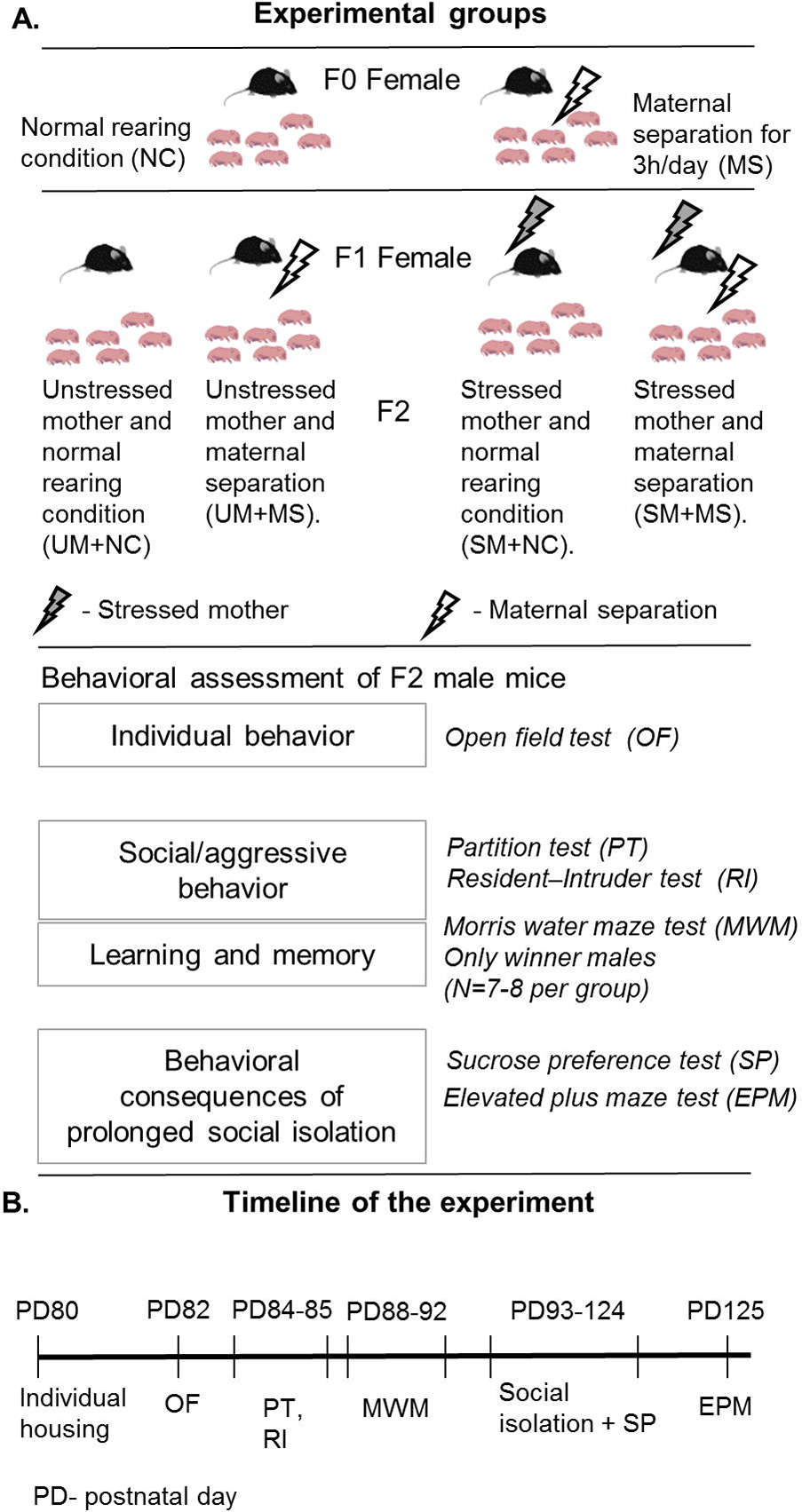
Experimental design and timeline. For detail explanations, see the text.

Only adult F2 males were used in behavioral testing. No more than two male offspring from each litter were used. Each group included 11-12 animals.

### 2.3. Behavioral tests

We used a battery of behavioral tests to assess the consequences of mothers’ past experience, pups’ stress and stresses in combination. Individual behaviors were evaluated in the open field test; aggressive behavior, in resident-intruder test; and cognitive abilities, by the Morris water maze test. Next, we investigated the influence of prolonged social isolation on the development of anxiety and depressive states in mice using the sucrose preference test and elevated plus maze.

Before the tests, mice were placed individually in experimental steel cages (14 × 28 × 10 cm) to abolish the group effects. Two days later, the behavioral tests were conducted in the following order: the open field, partition test, resident-intruder test, and Morris water maze (one test per day, the timeline is given in Figure 1). After that, all mice were exposed to social isolation for 32 days and then tested for sucrose preference and in the elevated plus maze test.

#### 2.3.1. Open field test

The open field (OF) consisted of a square arena (80 cm × 80 cm) with a white floor and 25-cm-high walls. The arena was brightly illuminated, had a central zone (40 cm × 40 cm) and a peripheral zone. Each mouse was placed individually in the center of the arena and the parameters of locomotor activity (distance traveled, mobility, average velocity and fast movements), exploratory activity (number of rearing, area covered) and anxiety (time spent in the central zone) were recorded over a 5-min test period. Area covered and number of rearing were scored using the EthoStudio software (Kulikov et al., 2005), the other parameters were scored using the EthoVision 10.0 software (Noldus Information Technology, Wageningen, the Netherlands). Fast movements were defined as any movements with a velocity higher than 10 cm/s. The open-field arena was thoroughly cleaned between animals.

#### 2.3.2. Resident-intruder test

To assess aggressive behavior, the experimental cage was bisected by a perforated transparent partition allowing the animals to see, hear, and smell each other but preventing any physical contact. Immediately, an unfamiliar male mouse of the same strain (an intruder, not significantly differing in weight any residents ± 0.3g) was placed behind the partition. After 5 min of habituation (partition test, see 2.3.3), the partition was removed to allow the resident and the intruder to interact for 10 min. After that, the partition was set up again and the intruder was left behind it for 24 h. On the next day, the repeated testing of partition behavior and resident-intruder confrontations was conducted. We used a two-day version of the test to assess behavioral changes following an overnight coexisting with the intruder. The social status in the model of coexisting with an intruder becomes clear on day 2 or 3 in confrontation (Bartolomucci et al., 2005), so the amount of time spend divided by the partition and having no physical contact may be important for developing a social coping strategy (Koolhaas, 1999). During the resident-intruder interactions, the following behavioral domains were analyzed: (1) direct attacks (biting, boxing, wrestling, tumbling the partner); (2) dominant behavior (chasing, aggressive grooming (keeping down the opponent, uprising front paws on the opponent)); (3) body sniffing (nose-to-whole body investigation of an area other than anogenital); and (4) anogenital sniffing (nose-to-anogenital area investigation). Latency to the first event, and total time of each behavioral domain were recorded. Behavioral data were collected using the free open-source software BORIS (Behavioral Observation Research Interactive Software) (Friard and Gamba, 2016).

#### 2.3.3. Partition test

To assess the reaction to another mouse (social behavior), the behavior of an experimental resident mouse near the partition bisecting the experimental cage was recorded for 5 minutes (Kudryavtseva et al., 2002, Kudryavtseva, 2003). The number of approaches to and the total time spent near the partition were scored. Each animal was tested twice with the same partner: on test day 1 immediately after placing the intruder behind the partition (before the first aggressive interaction) and on test day 2 after overnight cohabitation (before the second aggressive interaction).

#### 2.3.4. Morris water maze test

Spatial memory and learning were tested by using the Morris water maze paradigm (Morris, 1984). A circular pool (D=100 cm) was filled with water made opaque with the addition of powdered milk; the water temperature was maintained at a range of 22–24°C. The pool was divided into four quadrants with four starting locations (Target, Opposite, Sector 1 and Sector 2) marked by different visual cues on the pool wall. A hidden platform (d=8 cm) was located 1.5 cm below the water in the center of the Target sector. Mice were trained for 4 consecutive days and received 4 trials per day, each time randomly starting from a different starting point with a 15-sec interval between trials. After each trial, the mice were left on the hidden platform for 15 sec and then were given 15 sec for rest in the home cage before the next trial. To assess the learning ability, latency to find the platform in each of the 16 trials (4 trials/day × 4 days) was measured. Long-term spatial memory was assessed as latency to find the platform in each first trial on days 2-4, while short-term spatial memory was tested in a probe trial 15 min and 1 h after the last training trial on test day 4. For each probe trial, the hidden platform was removed, and the mice were placed into the pool for 60 sec, starting the probe trial from the Opposite sector. To assess spatial short-term memory, time spent in the Target sector and latency to find the previous location of the platform were measured. Parameters in the MWM test were scored using the EthoStudio software (Kulikov et al., 2005).

#### 2.3.5. Social isolation protocol

After the MWM test, the mice were individually housed in the experimental steel cages for 32 days with weekly cage cleaning.

#### 2.3.6. Sucrose preference test

For assessing stress-induced anhedonia, a sucrose preference test was conducted as previously described (Bondar et al., 2018a). The test consisted of a two-bottle choice and each mouse was allowed to choose between consuming water and a 1% solution of sucrose. The sucrose solution and water were provided on day 29 of social isolation. On the first day of exposure to the sucrose solution, the mice were allowed to adapt to the testing conditions, and the parameters of consumption were recorded on the second and third days of sucrose provision (measurement 1 and 2, respectively). The bottles were weighed to estimate the consumption of the liquids. The sucrose solution preference (the sucrose solution consumed as a percentage of the total amount of liquid intake) was measured.

#### 2.3.7. Elevated plus maze

The parameters of anxiety-like behavior as some of the consequences of prolonged social isolation (Fone and Porkess, 2008) were evaluated by the elevated plus maze test. All conditions and apparatus were as previously described (Bondar et al., 2018b). Briefly, to reduce the aversive conditions of the test, the experimental room was dimly lit. The parameters of anxiety-like behavior such as time spent in the open and closed arms as well as level of open arm area exploring (open arm area covered) were measured during a 5 min test period using the EthoStudio software (Kulikov et al., 2005).

### 2.4. Statistical Analysis

The normal distribution and homogeneity of variances were tested using Shapiro–Wilk’s and Levene’s tests, respectively, for all behavioral data. In general, the behavioral data were normally distributed and one-way ANOVA with the main factor being “type of stress” and the Fisher LSD test as a post hoc test were used to identify statistical differences between groups. The behavioral data from the resident-intruder test were not normally distributed, nonparametric tests were used. The behavioral data in this test was performed using Kruskal–Wallis test, with the type of stress as a factor. Pairwise comparisons were performed by the Mann–Whitney U test. The statistical analyses were conducted using the STATISTICA 8 software and the significance threshold for all behavioral parameters was set at *p* < 0 05.

## 3. Results

### 3.1. Body weight

Body weight was measured on PD14, 28 and 56. We found an influence of stress type on body weight on PD14 and PD28 (one-way ANOVA, PD 14: F(3,82)=3.87, p=0.012; PD28: F(3,69)=3.39, p=0.023). In SM+NC and UM+MS groups of males, the body weight on PD14 was reduced compared to the UM+NC group (p<0.05, Figure 2). It should be noted that SM-NC males, similar to UM-MS males, had a reduced body weight, although they were not physically separated from their mothers. Low weight in the mice of these groups appears to be due to impaired mother-offspring interaction. The SM+MS group demonstrated only a tendency to have a reduced body weight on PD14 (6.08 ± 0.08 vs 6.41±0.11 in the UM+NC group, p=0.11). Weight reduction in males is not a consequence of the change in the males-to-females ratio in the litter, because the effects on body weight in female offspring were similar (data not shown). At the same time, SM+MS males demonstrated increased body weights on PD28 compared with the UM-MS and UM-NC groups (p<0.05). These results may reflect increases in weight gain rates in SM+MS mice after maternal separation stress was eliminated.

**Figure 2.**
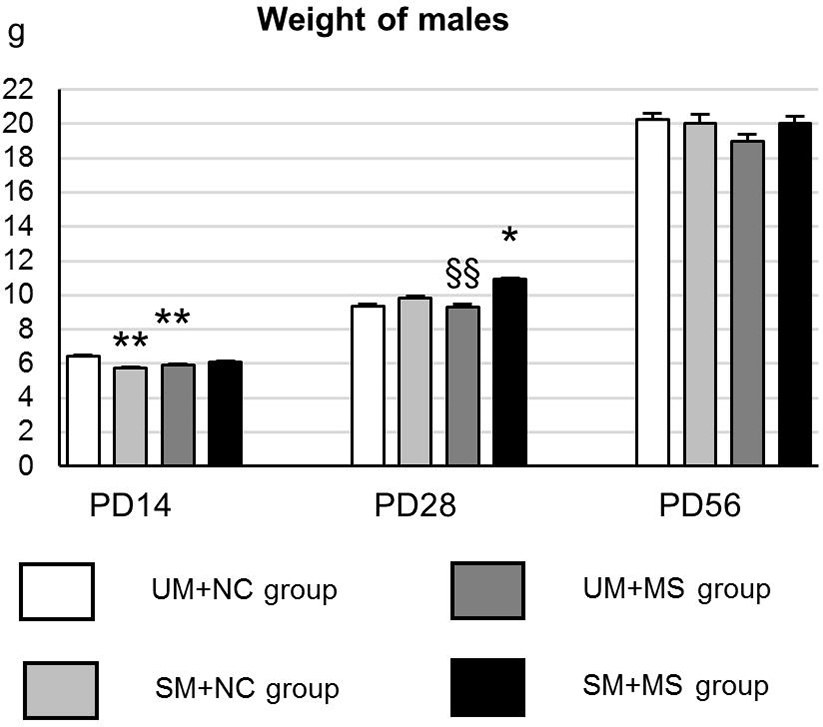
Body weight. Data are presented as mean ± SEM. * p<0.05, ** p<0.01 compared to the UM+NC group, §§ p<0.01 compared to the SM+MS group; Fisher’s LSD test as post hoc.

### 3.2. Mothers’ past stressful experience in the form of maternal separation in early life changes the social coping strategy, but not individual behavior

*Resident-intruder test.* Placing an unfamiliar partner to the cage, in which an experimental mouse had been residing for several days, led to aggressive confrontations. After two days of aggressive confrontations for 10 min each, we separated the mice into two groups: losers (those that had been defeated at least once) and winners (those that had not been defeated), the latter consisting of high-aggression males with a direct attack toward the enemy, and low-aggression males demonstrating only dominant behavior toward the partners.

Number of animals with different social statuses per group is shown in Figure 3B. The parameters of aggressive behavior in the animals are given in Figure 3C-G; no behavioral parameters of defeated animals are shown, for their number in the groups is low. On test day 1, all experimental groups had similar levels of latency to direct attack (Kruskal-Wallis test: H (3,31)=2.31, p=0.51), direct attack duration (H (3,28)=1.02, p=0.80), dominant behavior (H (3,28)=1.86, p=0.06), anogenital sniffing (H (3,28)=2.86, p=0.41) and sniffing (H (3,28)=6.77, p=0.08). Prolonged cohabitation (overnight) with intruders behind the partition led to increase aggression only in SM+MS males. Aggressive SM+MS males demonstrated decreased latency to direct attack (*U* = 9.0, *p* = 0.009) and increased direct attack duration (U=8.5, p=0.041) compared to test day 1. In addition, aggressive SM+MS males showed reduced levels of body and anogenital sniffing (U=0.0, p=0.002; U=3.0, p=0.006; and U=14.0, p=0.004, respectively) compared to test day 1. The other groups showed no significant changes in direct attack, although the animals showed reduced anogenital sniffing (p <0.05 for all groups) and dominant behavior (p <0.05 for UM+NC and UM+MS), but not sniffing compared to test day 1. Nevertheless, the behavior of aggressive males in the different groups did not differ significantly on test day 2. Increased aggression in SM+MS males on test day 2 is probably a feature of their social coping strategy. Thus, our results demonstrated that mothers’ stressful experience in combination with maternal separation in early life leads to a change in the social coping strategy – at least in aggressive males after prolonged cohabitation with intruders.

**Figure 3.**
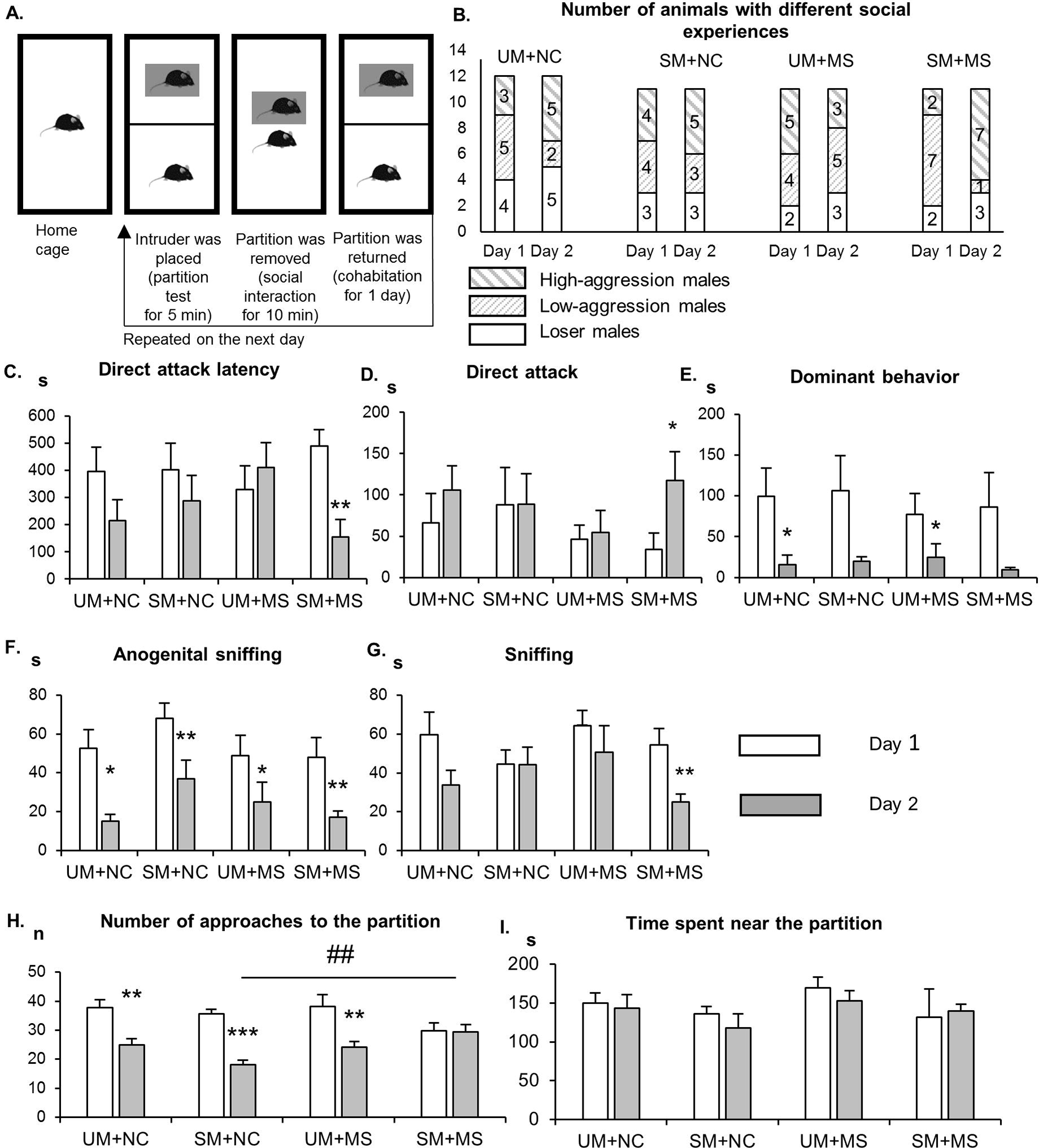
Behavior of F2 males in the resident-intruder and partition test. **A.** Experimental design of the resident-intruder test. **B.** Number of animals with different social experience. **C-G.** Behavior of aggressive males in agonistic confrontations. **H-I**. Behavior of aggressive males near the partition. Data are presented as mean ± SEM. * p <0.05, ** p <0.01, *** p <0.00 compared to test day 1. ## p<0.01 compared to the SM+NC group. Pairwise comparisons were performed by the Mann–Whitney U test.

### Open field test

We assessed the different parameters of individual behavior in the OF test. We did not find any significant differences between the groups (Figure 4). All groups demonstrated similar locomotor activity (distance traveled was more than 30 m and mobility was more than 50% of the test time), exploratory activity (number of rearing was more than 20 and the area covered was more than 70%) and anxiety (time spent in the center was more than 20%).

**Figure 4.**
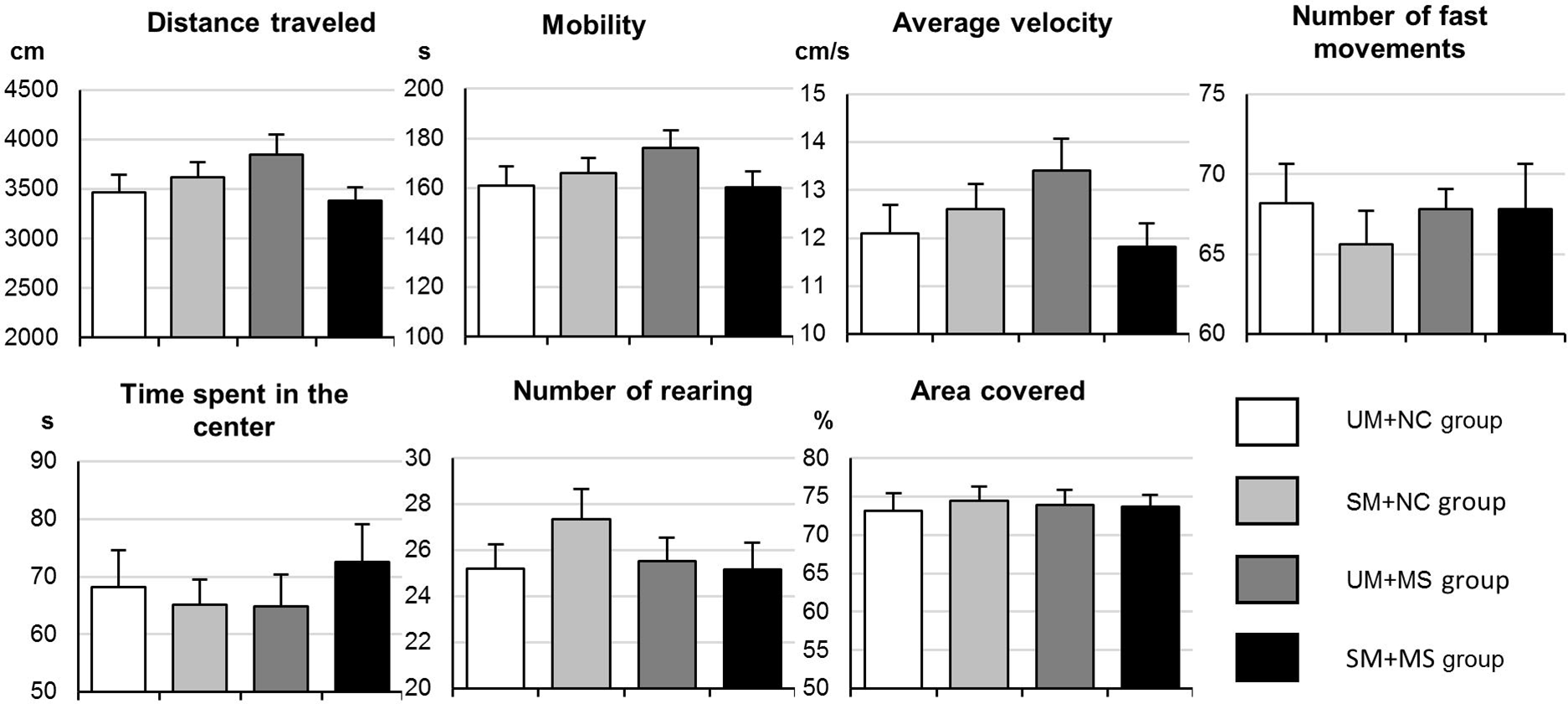
Behavior of F2 males in the open field test. Data are presented as mean ± SEM.

### Partition test

On test day 1, after placing an intruder behind the partition, we measured the reaction of the mice to new unfamiliar partners, which may reflect the level of aggressive motivation (Kudryavtseva et al., 2002, Kudryavtseva, 2003). There were no significant differences between the groups; in all groups, mice spent more than 50% of time near the partition and made similar numbers of approaches to the partition (Kruskal-Wallis test: H (3,45) = 3.03, p = 0.390 and H (3,45) = 2.27, p = 0.520, data not presented). We separately considered the partition behavior of males that displayed aggressive behavior (Figure 3H-I). We found that past stress experience influences the number of approaches to the partition on test day 2 (Kruskal-Wallis test: H (3, 31) = 11.83 p =0.008), but not the time spent near the partition (H (3, 31) = 3.42 p =0.33). Aggressive SM+MS males made more approaches to the partition compared to SM+NC males (p = 0.005). A comparison of the behavior of aggressive males on test days 1 and 2 showed that the aggressive males of all groups except SM+MS made less approaches to the partition on test day 2 than on test day 1 (p <0.01); while aggressive SM+MS males demonstrated a similar number of approaches on test day 1 and 2 (U = 21.5, p = 0.747). These results may indicate the lack of reduction in aggressive motivation in aggressive SM + MS males following prolonged cohabitation with intruders.

## 3.3. Aggressive males produced by mothers stressed by maternal separation in their early life have improved memory

### Morris water maze test

Learning was evaluated by latency to find the platform in each trial. All groups demonstrated a typical learning curve with a decrease of escape latency across trials (Figure 5A). We found an effect of past experience on latency to find the platform on test day 3 (repeated one-way ANOVA, F(3,26)=3.38, p=0.033). Aggressive SM+NC males showed an increased time to find the platform (p<0.05) compared to males in the other groups. (Latency to find the platform in the first trial on day 2-4 was considered a parameter of long-term spatial memory) (Figure 5B). We found significant effects of past experience on latency to find the platform in first trial only on test day 2 (repeated ANOVA, day 2: F(3, 26)=8.49, p<0.001; day 3: F(3,26)=1.29, p=0.300; day 4: F(3,26)=2.04, p=0.133). Aggressive SM+MS males were quicker in locating the platform in the first trial on training day 2 than the males in other groups. However, the SM+MS group did not change this parameter throughout the training days, while the aggressive males in the SM+NC, NC+MS and NC+NC groups demonstrated decreases in latency to find the platform in the first trial on test day 4 compared with test day 2 (p<0.001, p=0.017 and p=0.092 respectively). Thus, the aggressive SM+MS males have a better memory for the location of the platform after the first day of the test, while the other groups gradually memorize visual cues.

**Figure 5.**
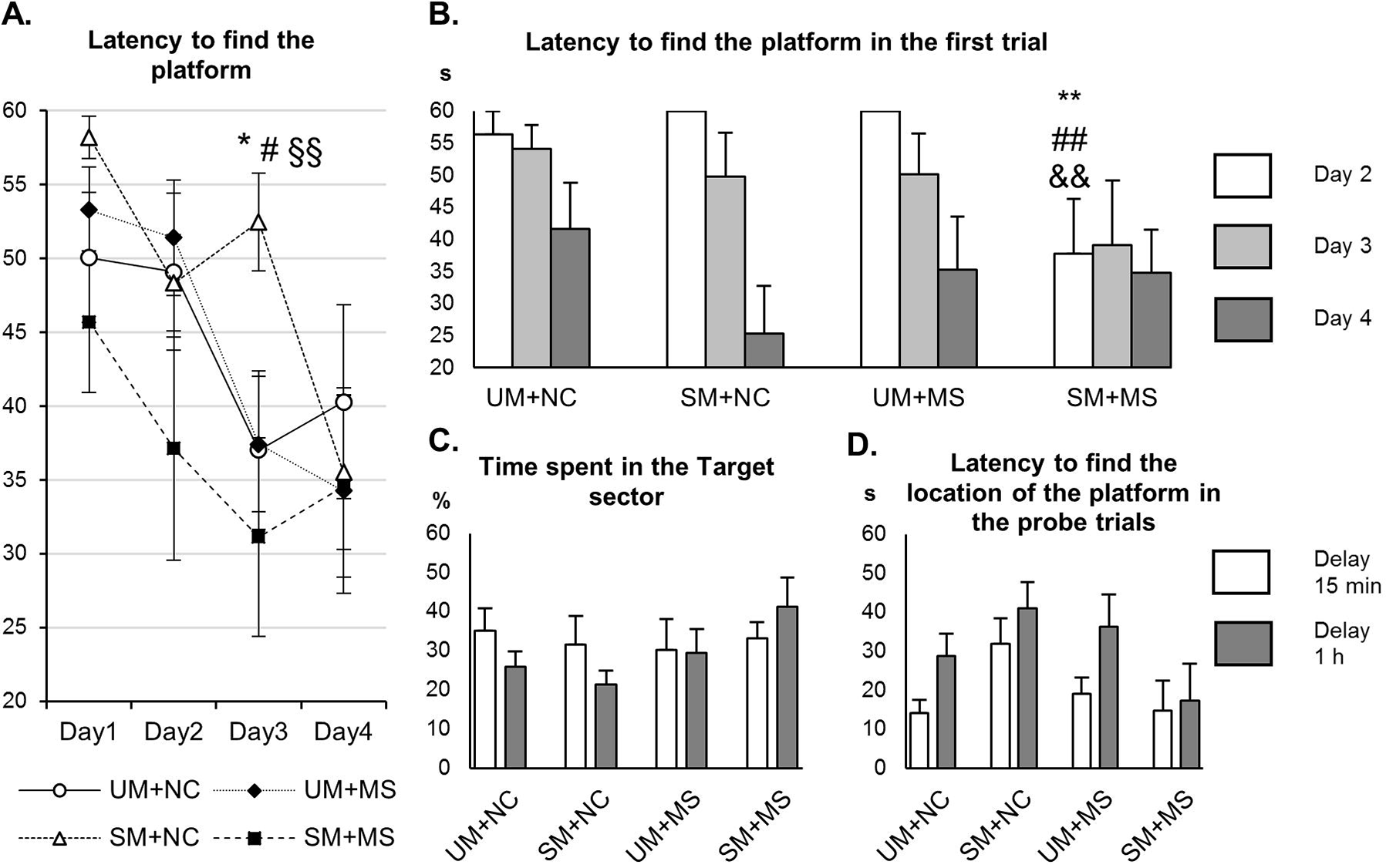
Behavior of aggressive males in the MWM test. **A.** Latency to find the platform during each training trial **B.** Latency to find the platform in the first trial on training days 2-4. Spatial long-term memory on training days 2-4. **C-D.** Time spent in the Target sector and latency to find the previous location of the platform were measured in a probe trial 15 min or 1 h after the last training trial as parameters of spatial short-term memory. Data are presented as mean ± SEM. * p <0.05; ** p <0.01 compared to the UM+NC group, # p <0.05, ## p <0.01 compared to the UM+MS group; && p <0.01 compared to the SM+NC group; §§ p <0.01 compared to the SM+MS group; Fisher’s LSD test as post hoc.

To assess short-term memory, the mice were re-exposed to the Morris water maze 15 min and 1 h after the last training trial on test day 4.

Neither delay in re-exposure revealed any differences in memory retrieval between the groups: neither time spent in the Target sector nor latency to find the previous location of the platform (Figure 5C) differed between the groups. Admittedly, there was a subtle tendency for effects of past stressful experience following re-exposure after 1 h (one-way ANOVA, F(3.26)=2.05 p=0.131) on the time spent in the Target sector. SM-MS mice had a slightly higher time spent in the Target sector compared with the other groups.

All groups spent over 30% of time in the Target sector after 15 minutes’ delay in re-exposure, while after 1 hour’s delay this parameter was decreased in the winner males of all groups except SM-MS (40%). In general, our results demonstrate learning impairments in aggressive SM-NC males, while SM-MS males have signs of improved long-term spatial memory.

## 3.4. Past stress has no effect on the consequences of prolonged social isolation

To assess behavioral changes occurring on the background of social interaction deficits, we exposed males to prolonged social isolation for 32 days and assessed their states using tests for anxiety and anhedonia.

### Sucrose preference test

We did not find any significant differences in sucrose preference between the groups on test days 1 or 2 (Figure 6A). Aggressive males in all groups demonstrated sucrose preference over water consumption (sucrose solution consumption was more than 70% of all liquid intake) as unstressed animals (Bondar et al., 2018a).

**Figure 6.**
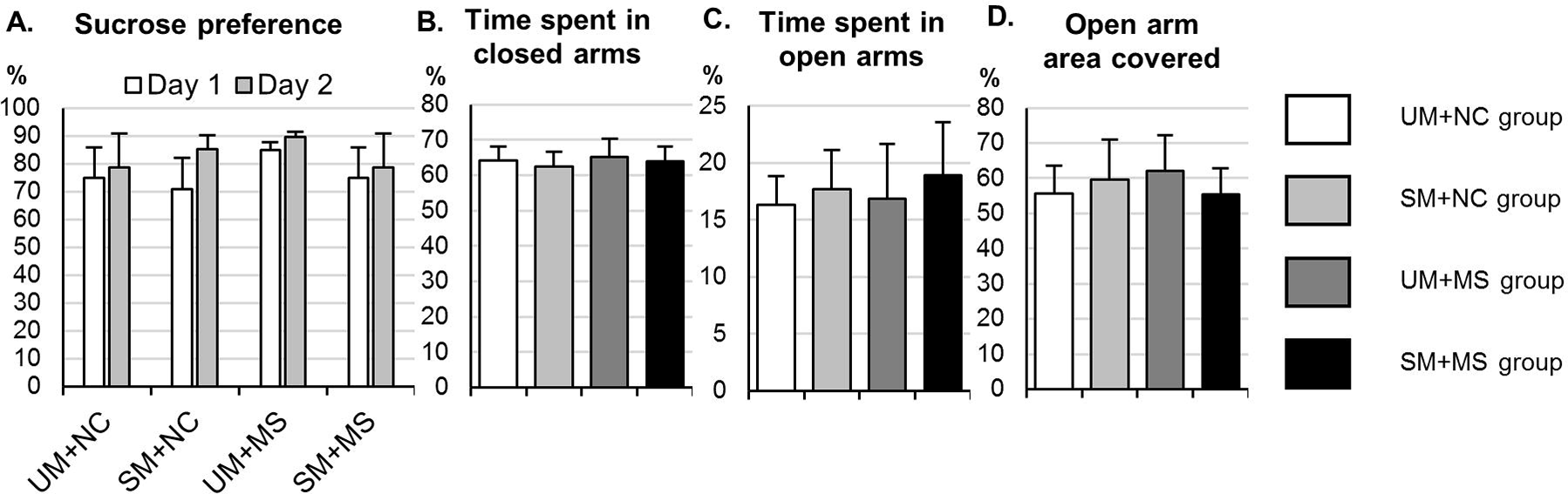
Behavior of aggressive F2 males after prolonged social isolation. **A.** Sucrose preference test **B-D.** Plus maze behavior. Data are presented as mean ± SEM

### Elevated plus maze test

Anxiety evaluated in the elevated plus maze did not differ between the groups (Figure 6B). Aggressive males in all behavioral groups had similar times spent in closed arms (more 60%), open arms (more 15%) and open arm area covered (more 50%).

## 4. Discussion

Mother-offspring interactions during early postnatal times plays a key role in neuronal development and shaping adult behavior (Cirulli et al., 2003). Disruptions in these interactions caused both by physical separation of pups from their mothers and by stress experienced by the mothers during and before lactation can lead to changes in the behavior of their adult offspring (Johnson et al., 2011, Kosten et al., 2012, Babb et al., 2014, Steenwyk et al., 2018). In the present study, we found that stresses applied in combination (stressed mother + maternal separation, SM+MS) have no effect on aggressiveness, but leads to changes in the social coping strategy on the background of prolonged agonistic social contacts. At the same time, neither mice raised by a stressed dam without early-life stress (SM+NC) nor mice with a history of early-life stress alone (UM+MS) had difference from controls in social behavior; they had similar social coping strategies on the background of agonistic social contact. Additionally, aggressive SM+MS males tend to have improved long-term spatial memory, while SM+NC males showed signs of impaired learning in the MWM. We found no differences in the parameters of individual behavior or in behavior after prolonged social isolation. Thus, our findings demonstrated that the mothers’ past experience in their early life plays an important role in shaping the behavioral phenotype of their adult offspring.

The effects of early postnatal stress on social behavior and aggression in rodents remains poorly studied and the results obtained to date are often inconsistent. Studies in rodents demonstrate that prolonged maternal separation in early life has no effect on the duration of social contacts in juvenile or adult male rats (Hulshof et al., 2011, Holland et al., 2014, Farrell et al., 2016) or mice (Franklin et al., 2011, Tsuda et al., 2011). However, some researchers find that maternal separation in early life may lead to increases (Kundakovic et al., 2013) or to decreases (Niwa et al., 2011) in social behavior in adolescent mice. A similar inconsistency was shown for offensive play-fighting and aggressive behavior in juvenile and adolescent rodents. Play-fighting is an important behavior in adolescent males and is associated with the level of aggression in adulthood (Vanderschuren et al., 1997). Some researchers find decreases in play-fighting behavior and aggression in adolescent mice and rats (Muhammad and Kolb, 2011, Tsuda et al., 2011), while others observe increased aggression (Veenema and Neumann, 2009, Shin et al., 2016). Works on adult rodents showed that prolonged maternal separation leads to increased aggression in rats and decreased aggression in mice (Veenema et al., 2006, Veenema et al., 2007). These inconsistent results may be due to methodological differences (the number of test days) and the use of different strains of mice as intruders. Running the test for 2 days including cohabitation with an intruder allowed us to assess not only the level of social motivation, but also the strategy of social habituation (Koolhaas et al., 1999, Bartolomucci et al., 2004). Indeed, we did not find any significant differences in aggression or social behavior between the groups on the first or second day of aggressive interactions; however, aggressive SM+MS males became even more aggressive after the first day of cohabitation with intruders; additionally, that was the only group that did not decrease aggressive motivation, no matter how long cohabitation lasted. Thus, these results are thought to provide evidence of changes in the social coping strategy, the changes being associated with changes in aggressive motivation in aggressive SM+MS mice due to lasting contacts with conspecific intruders.

Surprisingly, in addition to having changes in the social coping strategy, the aggressive SM+MS males were quicker to find the platform in the first trial of MWM test day 2 than any other group, suggesting a tendency toward enhanced spatial long-term memory. At the same time, aggressive SM+NC males tended to be slower to learn on MWM test day 3. Long-term spatial memory strongly depends on the circulating levels of glucocorticoid hormones during acquisition and consolidation phase (Lupien and Lepage, 2001, Kim and Diamond, 2002, Conrad, 2005, Taylor et al., 2015). When tasks are minimally aversive, glucocorticoid levels behave as a non-linear function of hippocampal-dependent memory; during acquisition and consolidation, optimal performance is reached at low-to-moderate glucocorticoid levels; and impaired memory is observed occurs at very low or very high glucocorticoid levels (Lupien and Lepage, 2001, Kim and Diamond, 2002). In contrast, highly aversive paradigms during acquisition or exposure to pre-training stress may lead to a linear association between glucocorticoid levels and memory consolidation (Conrad, 2005, Taylor et al., 2015). Considering this, it is possible that the observed tendency towards enhanced long-term spatial memory in aggressive SM+MS males is associated with changes in HPA axis reactivity to stress produced by the MWM (Harrison et al., 2009) or to past stressful experience gained in the resident-intruder test (Bhatnagar et al., 2006), or to both in combination. These factors could be the reason for the absence of changes in the spatial memory of aggressive males in the UM+MS group. Rodent studies, as our original works, demonstrate that prolonged maternal separation paradigms usually lead to cognitive and learning impairments in rats (Kosten et al., 2012) and mice (Mehta and Schmauss, 2011, Reshetnikov et al., 2018a, Reshetnikov et al., 2018b).

The hypothalamic–pituitary–adrenal (HPA) axis is the most important effector of the stress response; many investigators suggest that early-life stress may be a reason behind long-term changes in HPA functionality (Heim et al., 2004, Fumagalli et al., 2007, Juruena, 2014). Human studies demonstrated that HPA axis and autonomic nervous system hyperreactivity may— presumably due to CRF hypersecretion and reduced inhibitory feedback—be a persistent consequence of childhood abuse that may contribute to the diathesis for adulthood psychopathology (Heim et al., 2008, Von Werne Baes et al., 2012). Adult rats exposed to maternal separation show significantly enhanced pituitary-adrenal responses to acute stress which may result from blunted negative feedback due to reduction in GR binding in the hippocampus and enhanced expression of Crh mRNA in the hypothalamus (Plotsky and Meaney, 1993, Liu et al., 2000, Ladd et al., 2004). In support of these data, our previous study demonstrated that MS resulted in reduced hippocampal Crhr1 mRNA, an increased MR/GR mRNA ratio in the hippocampus and hypothalamus and increased Avp mRNA in the hypothalamus (Reshetnikov et al., 2018b). Thus, our previous study showed that prolonged separation of pups from their mothers leads to pronounced changes in HPA-associated gene expression.

Finally, an elegant work by Weaver and the co-workers (2004) showed that offspring receiving high levels of active maternal care had reduced DNA methylation and increased expression of GR in the hippocampus and, therefore, enhanced negative feedback sensitivity to glucocorticoids. Thus, changes in mother-offspring interactions leading to changes in active maternal care given to the offspring, as early-life stress, may lead to a misbalance between excitatory and inhibitory activity input in response to stress. Considering this, we can hypothesize that the changes we found in social behavior may long-term spatial memory in the aggressive males of the SM+MS group and assessed in highly stressful stress test conditions might be caused by enhanced stress reactivity associated with changes in the HPA axis functionality. It is possible that differences may be both adaptive and non-adaptive.

In this study, we did not find any behavioral differences in individual behavior between the groups. Data on the effects of early-life stress on open-field behavior in adulthood are often inconsistent. A high percentage of studies reported that prolonged maternal separation does not significantly change open-field behavior in male mice (Loi et al., 2017). At the same time, some studies in mice, including our previous work, found that prolonged maternal separation leads to reduced locomotor activity (Varghese et al., 2006, Bondar et al., 2018b). Tractenberg and the co-workers (2016) reported that differences in the results of studies in mice may be due to methodological differences in the procedure of maternal separation: the latter differences may have effects on the level of stress received by pups. For example, maternal separation of pups that had been housed individually and pups that had been kept in litters results in quite different levels of tactile stimulation, which are essential for adequate development (Kaffman and Meaney, 2007). Considering this fact, we propose that variations in the maternal separation procedure (individual housing in this study or keeping in litters in our previous report (Bondar et al., 2018b)) could have different implications for the parameters of their individual behavior in adulthood.

In the final part, the behavioral consequences of prolonged social isolation in aggressive males were measured. Although prolonged social isolation in rodents normally produces different types of behavioral dysfunctions such as depressive‐ and anxiolytic-like behaviors (Fone and Porkess, 2008, Chang et al., 2015), we did not find any behavioral differences in anxiety or depressive-like behavior between groups. Lack of differences may be regarded as evidence of both insufficient duration of social isolation for developing depressive‐ and anxiolytic-like behaviors and insufficiently aversive test conditions, in which these behavioral parameters were measured. Another factor with implications for behavioral effects of prolonged social isolation could be stressful experience gathered in previous tests. Unfortunately, a limited body of data prevent us from identification of the true reason and all our hypothesizing remains to be speculative.

Overall, our findings suggest that mothers’ past experience combined with maternal separation in early life (SM+MS) has the strongest effect on the behavior of adult offspring. These findings, together with analysis of behavior in aggressive males raised by stressed dams (SM+NC), allowed us to come to the conclusion that mothers’ past experience have a significant effect on offspring behavior. We suggest that the observed results are most likely due to disruptions in mother-offspring interactions. Impaired maternal care in MS mothers toward their pups may be regarded as one of the possible delayed effects of these mothers’ early-life stress. In rodents, licking and other tactile stimulation of pups play the most essential role in furthering offspring behavior (Rosenblatt, 1967, Numan et al., 1990, Shoji and Kato, 2009). Studies in rats indicating that the level of maternal care received by offspring in their early life have effects on their behavior in adulthood (Francis et al., 1999, Champagne et al., 2003, Jensen Pena and Champagne, 2013, Pan et al., 2014). These studies revealed that female offspring of the mothers with low levels of maternal care show decreased locomotor activity and increased anxiety in their adulthood (Pedersen et al., 2011, Jensen Pena and Champagne, 2013, Pan et al., 2014). It should be noted that adult female rats that were exposed to repeated periods of maternal separation or chronic social stress in their early life exhibit reduced maternal licking and crouching over their pups (Lovic et al., 2001, Carini and Nephew, 2013). The relevant human studies also showed that the maternal adverse childhood experiences such as maltreatment, abuse and neglect affect the development of their infant (Berthelot et al., 2015, Racine et al., 2018). At the same time, the study in mice demonstrated a slight effect of early-life stress in female offspring on their maternal care in adulthood (Weiss et al., 2011). However, regardless of the changes in the level of maternal care, the stress experienced by mothers in their early life led to a change in the adult behavior of their own progeny (Weiss et al., 2011, Babb et al., 2014, Carini et al., 2013), which, in turn, confirms the effects of the mothers’ past experience on the development of their offspring.

In summary, to the best of our knowledge, ours is the first study in mice that compares the effects of the mothers’ past stress experience on F1 females, the effects of prolonged maternal separation on F2 males in early life or the effects of both stresses in combination on the behavior of male mice in adulthood. We found that the combination of stresses (stressed mother + maternal separation, SM+MS) led to the most pronounced effects on offspring behavior. Our findings bring us to the conclusion that past stressful experience in a mother may modulate the effects of early postnatal stress in her offspring and that this modulating action is probably mediated by changes in mother-offspring interactions. However, the molecular background these effects remain unclear. Future research is required to identify possible molecular mechanisms and pathways by which mothers’ past experience impact their adult offspring behavior.

